# Expression of β-catenin, LEF-1, MMP7 and SFRP5 associated with poor outcome in pancreatic cancer

**DOI:** 10.1101/2025.06.11.659232

**Authors:** Panpan Kong, Feng He, Huifang Liu, Dong Yan

## Abstract

Pancreatic cancer is highly aggressive and lethal malignancies worldwide. β-catenin, LEF-1, MMP7 and SFRP5 are key regulatory proteins in the Wnt signaling pathway. It has been confirmed that Wnt signaling pathway is involved in the biological regulation of pancreatic cancer cells and plays an important role. However, its effect on the malignant phenotypes of pancreatic cancer cell and the corresponding molecular details remains unknown. In the current study, the expression of β-catenin, LEF-1 and MMP7 was up-regulated and the expression of SFRP5 was down-regulated in pancreatic cancer cell lines and tissues by RNA sequencing, quantitative PCR (qPCR) and immunohistochemical analysis. β-catenin, LEF-1 and MMP7 positive expression correlates with poor prognosis in PDAC patients, and SFRP5 positive expression correlates with longer survival times. Subsequently r software was used to establish a rosette model to predict the prognosis of pancreatic cancer according to the results of multi-factor analysis. After statistical analysis, age, CA-199, β-catenin, MMP7 and SFRP5 could serve as promising biomarkers for predicting prognosis in PDAC patients. This study also suggests that survival probability of patients can be reliably predicted by a nomogram-based method.

## Introduction

Pancreatic cancer is one of the tumors with the highest mortality rate among solid tumors. Distant metastasis through nerve, vascular and lymphatic invasion pathways greatly reduces the survival rate of patients and has become the third major cause of cancer-related death worldwide[1, 2]. According to 2022 Global Cancer Statistics, the number of new cases of pancreatic cancer reach 510,566 of which approximately 467,005 people died of this cancer, the mortality rate is almost similar to its incidence rate[3]. Although many treatments such as surgery, chemotherapy, radiotherapy and interventional therapy have been applied in the treatment of pancreatic cancer, the clinical efficacy is remains unsatisfactory. Radical resection is considered the only curative approach for pancreatic cancer[4, 5], but surgical resection rates have been reported to be extremely low (about 15%-20%), even after a potentially radical resection, up to 80% of patients eventually relapse within 2 years[6, 7]. Thus, the treatment of pancreatic cancer has become one of the world’s unsolved dilemma. To date, despite the considerable scientific progress achieved about signaling pathways along with its molecular mechanism studies, but these progresses have not yet translated into more effective therapeutic strategy for PDAC patients[8].

As a key regulatory network for embryonic development and adult tissue homeostasis, Wnt signaling pathways are widely studied due to their important roles in cell communication, cell regulation, and cancer occurrence[9]. The abnormal activation or disruption of Wnt signaling pathways are closely linked to several different diseases, such as cancer, alzheimer’s disease, osteoporosis, diabetes, etc.[10-12], and are potential targets for pharmacological therapy, but the exact mechanism behind this regulation remains fully elucidated.

In the preliminary experiment, we obtained the information of all the members of the WNT signaling pathway by searching the BioCarta Pathways database. In addition, we screened the differentially expression genes in PDAC tissue through the gene chips, and performed GO analysis and Pathway analysis. Based on the results of the chip analysis and BioCarta Pathways database, the members of the WNT signaling pathway with differential expression were screened out. The analysis results showed that compared with adjacent tissues in PDAC tissues, WNT signaling pathway members β-catenin, LEF1, MMP7, and SFRP5 all had differentially expressions.

Therefore, we aim to evaluate the long-term survival outcomes and influence factors of PDAC patients after radical resection, meanwhile, to explore the clinical significance of β-catenin, LEF-1, MMP7 and SFRP5 in PDAC. At the same time, a lirographic prognostic prediction system that integrates the anatomical and biological characteristics of tumor and is clinically easy to use was established, and the consistency test of predicted value and actual value was carried out with high accuracy, which is expected to be used as the prediction of overall survival time of patients.

## Patients and Methods

### Patient data and specimens

45 PDAC tissue specimens and corresponding adjacent normal tissues located >2 cm from the edge of the cancer tissue were obtained from patients who underwent radical pancreatoduodenectomy at the Affiliated Tumor Hospital of Xinjiang Medical University, Urumqi, Xinjiang Uygur Autonomous Region from June 2012 to December 2013. All patients did not receive chemotherapy, radiotherapy, or immunotherapy prior to surgery. The clinical data of patients were collected, including age, gender, body mass index (BMI), diabetes mellitus, total bilirubin, bilirubin direct, serum tumor markers, tumor characteristics, and valid survival data. The data that support the findings of this study are available via e-mail from the corresponding author upon reasonable request.

### Materials and reagents

PANC-1, HPDE6 were purchased from Bei Na Chuang Lian Biotechnology Institute, rabbit monoclonal antibody against human β-catenin , rabbit monoclonal antibody against human LEF-1,rabbit monoclonal antibody against human MMP7 and rabbit monoclonal antibody against human SFRP5 were from Zhongshan Goldenbridge Biotechnology, SP immunohistochemistry kit were from Fuzhou Maixin Biotech Co., Ltd..

### Quantitative real-time polymerase chain reaction (QRT-PCR)

Total RNA was extracted from cultured cells using the Trizol Reagent (Invitrogen, Carlsbad, CA, USA). The RNA was reverse-transcribed into cDNA using TransScript One-Step gDNA Removal and cDNA Synthesis SuperMix (Transgen, Beijing, China) according to the manufacturer’s instructions. QRT-PCR was performed to detect the expression level of mRNA. The primers were as follows (5’-3’): β-catenin: AAAGCGGCTGTTAGTCACTGG (forward), CGAGTCATTGCATACTGTCCAT (reverse); LEF-1: GCATCAGGTACAGGTCCAAGA (forward), TGTTCCTTTGGGGTCGACTG (reverse); MMP7: CATGATTGGCTTTGCGCGAG (forward), AGACTGCTACCATCCGTCCA (reverse); SFRP5: CTGTACGCGTCATCCTAGCC (forward), CGGACCAGAAGGGGGTCTAT (reverse); β-actin: CATGTACGTTGCTATCCAGGC (forward), CTCCTTAATGTCACGCACGAT (reverse). β-actin was used for internal control. The data were analyzed using the 2^-ΔΔCt^ method.

### Immunohistochemistry

Paraffin sections of tissue specimens were 5μm thick and stained with the SP kit according to the routine immunohistochemical procedure. Phosphate-buffered saline (PBS) was used as negative controls. The staining score is as follows: 0, no staining; 1+, minimum staining; 2+, moderate to strong staining of at least 20% of cells; 3+, strong staining of at least 50% of cells. 0 or 1+ stained cases were classified as negative, and those with 2+ or 3+ staining cases were considered positive.

### Statistical analysis

Continuous variables are presented as the mean ± standard deviation (SD), categorical variables as percentages (%) and counts. Categorical variables were compared using the Chi-square test and Fisher’s exact test. The Spearman rank correlation coefficient was used to analysis the correlation of protein expression. Overall survival (OS) curves were plotted by using the Kaplan–Meier method and compared by using log-rank test. Univariate and multivariate analysis for survival was performed using the Cox proportional hazard regression model. *P*-value <0.05 was considered statistically significant. Statistical analyses were performed using SPSS 26.0 software (IBM Corp., Armonk, NY, USA) and GraphPad Prism 8.0 software (GraphPad Software Inc., CA, USA).

## Results

### The mRNA expression levels of β-catenin, LEF-1, MMP7 and SFRP5 in PDAC cell lines

QRT-PCR analysis showed that except SFRP5, the mRNA expression levels of β-catenin, LEF-1 and MMP7 were significantly increased in PDAC cells (PANC-1) compared to normal pancreatic cells (HPDE6) (*P*<0.01, Figure 1).

**Figure 1.**
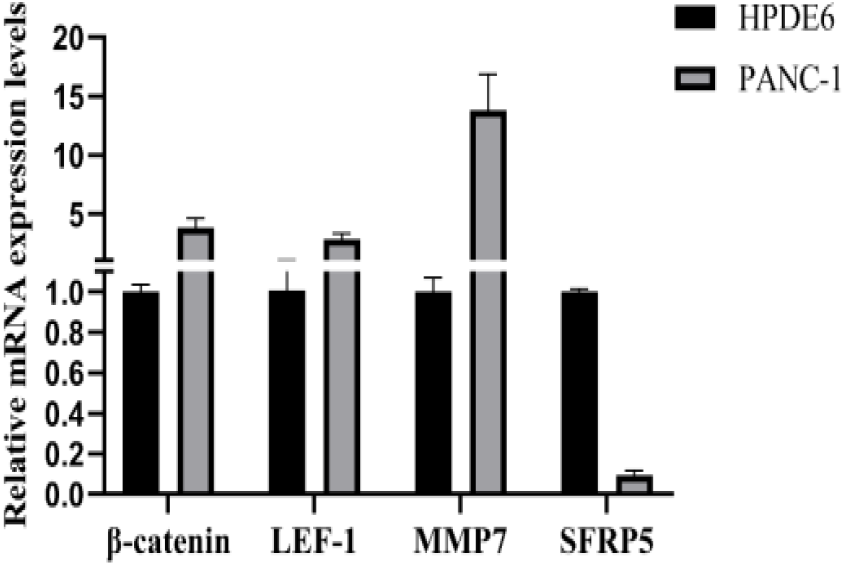
The mRNA relative expression of β-catenin, LEF -1, MMP7 and SFRP5 in PDAC and normal pancreatic cell lines.

### The expression of β-catenin, LEF-1, MMP7 and SFRP5 in PDAC tissues

As a result of immunohistochemical evaluation of the tissue specimens (Figure 2), In PDAC tumor tissues, β-catenin (p=0.005), LEF-1 (p=0.000), MMP7 (p=0.023) positive expression rates were significantly higher and SFRP5 (p=0.017) was lower than in adjacent normal tissues (Table 1). Negative correlation was found between the expression of LEF-1 and SFRP5. (Table 2)

**Table 1.** Expression of β-catenin, LEF-1, MMP7 and SFRP5 in PDAC and adjacent normal tissues.

|  | **Tumor(n=45)** | **Peritumor(n=45)** | **p-value** |
| --- | --- | --- | --- |
| **β-catenin** |  |  | 0.005** |
| Positive (%) | 31(68.9) | 15(33.3) |  |
| Negative (%) | 14(31.1) | 30(66.7) |  |
| **LEF-1** |  |  | 0.000*** |
| Positive (%) | 33(73.3) | 12(26.7) |  |
| Negative (%) | 12(26.7) | 33(73.3) |  |
| **MMP7** |  |  |  |
| Positive (%) | 26(57.7) | 14(31.1) | 0.023* |
| Negative (%) | 19(42.2) | 31(68.9) |  |
| **SFRP5** |  |  | 0.017* |
| Positive (%) | 26(57.7) | 34(75.6) |  |
| Negative (%) | 19(42.2) | 11(24.4) |  |

\* $P<0.05$ , \*\* $P<0.01$ , \*\*\* $P<0.001$ .

**Table 2.** Correlation between the expression of β-catenin, LEF-1, MMP7 and SFRP5 in PDAC tissues.

|  |  | **β-catenin** | **LEF-1** | **MMP7** | **SFRP5** |
| --- | --- | --- | --- | --- | --- |
| **β-catenin** | r | - | 0.346 | 0.009 | -0.203 |
|  | p-value | - | 0.103 | 0.955 | 0.181 |
| **LEF-1** | r | 0.346 | - | 0.095 | -0.298 |
|  | p-value | 0.103 | - | 0.535 | 0.046* |
| **MMP7** | r | 0.009 | 0.095 | - | -0.180 |
|  | p-value | 0.955 | 0.535 | - | 0.236 |
| **SFRP5** | r | -0.203 | -0.298 | -0.180 | - |
|  | p-value | 0.181 | 0.046* | 0.236 | - |

\* $P<0.05$ , \*\* $P<0.01$ , \*\*\* $P<0.001$ .

**Figure 2.**
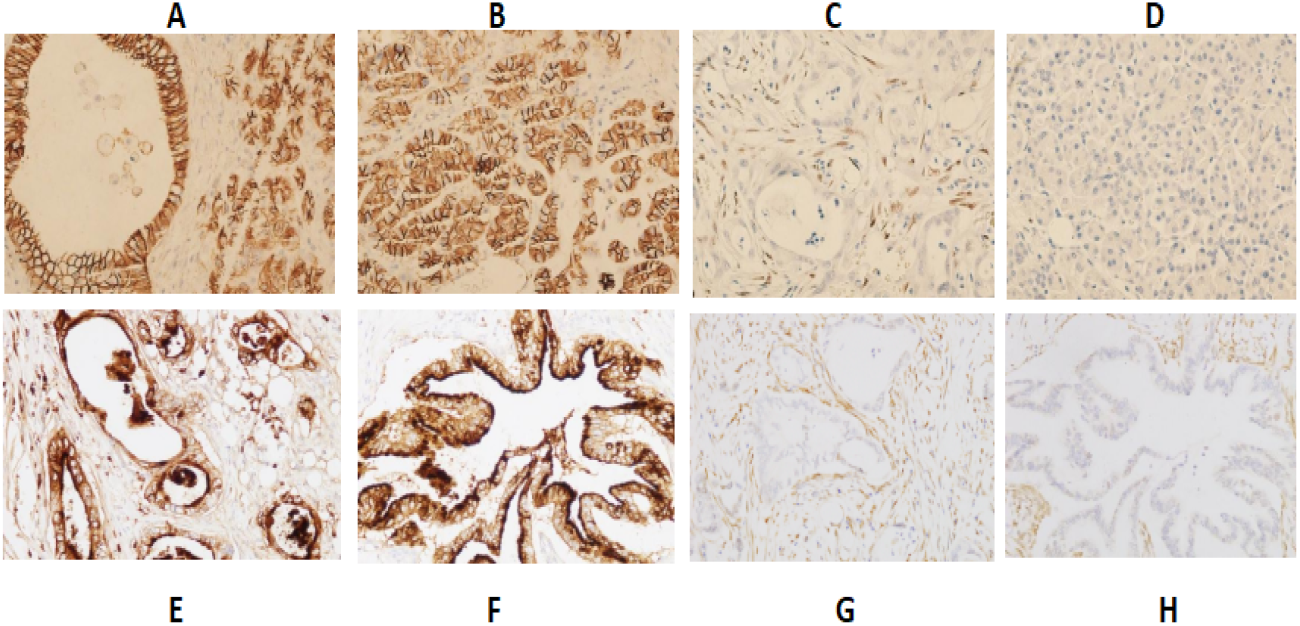
Expression of β-catenin LEF-1,MMP7 and SFRP5 in PDAC and adjacent normal tissues (SP × 400).(A) Expression of β-catenin in PDAC tissues. (B) Expression of β-catenin in adjacent tissues. (C) Expression of LEF-1 in PDAC tissues. (D) Expression of LEF-1 in adjacent tissues. (E) Expression of MMP7 in PDAC tissues. (F) Expression of MMP7 in adjacent tissues. (G) Expression of SFRP5 in PDAC tissues. (H) Expression of SFRP5 in adjacent tissues.

### Relationship between β-catenin/LEF-1/MMP7/SFRP5 and clinicopathological characteristics in PDAC patients

The clinicopathological characteristics of all patients and comparative analysis results between the β-catenin/LEF-1/MMP7/SFRP5 negative group and the β-catenin/LEF-1/MMP7/SFRP5 positive group were presented (Table 3 and Table 4). We found that β-catenin expression was related to tumor size (*P*=0.035); LEF-1 expression was related to serum CA19-9 level (*P*=0.001) and lymph node metastasis (P=0.041); SFRP5 expression was related to tumor differentiation (*P*=0.004).

**Table 3.**
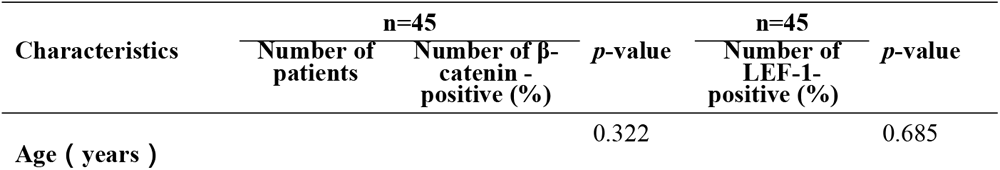
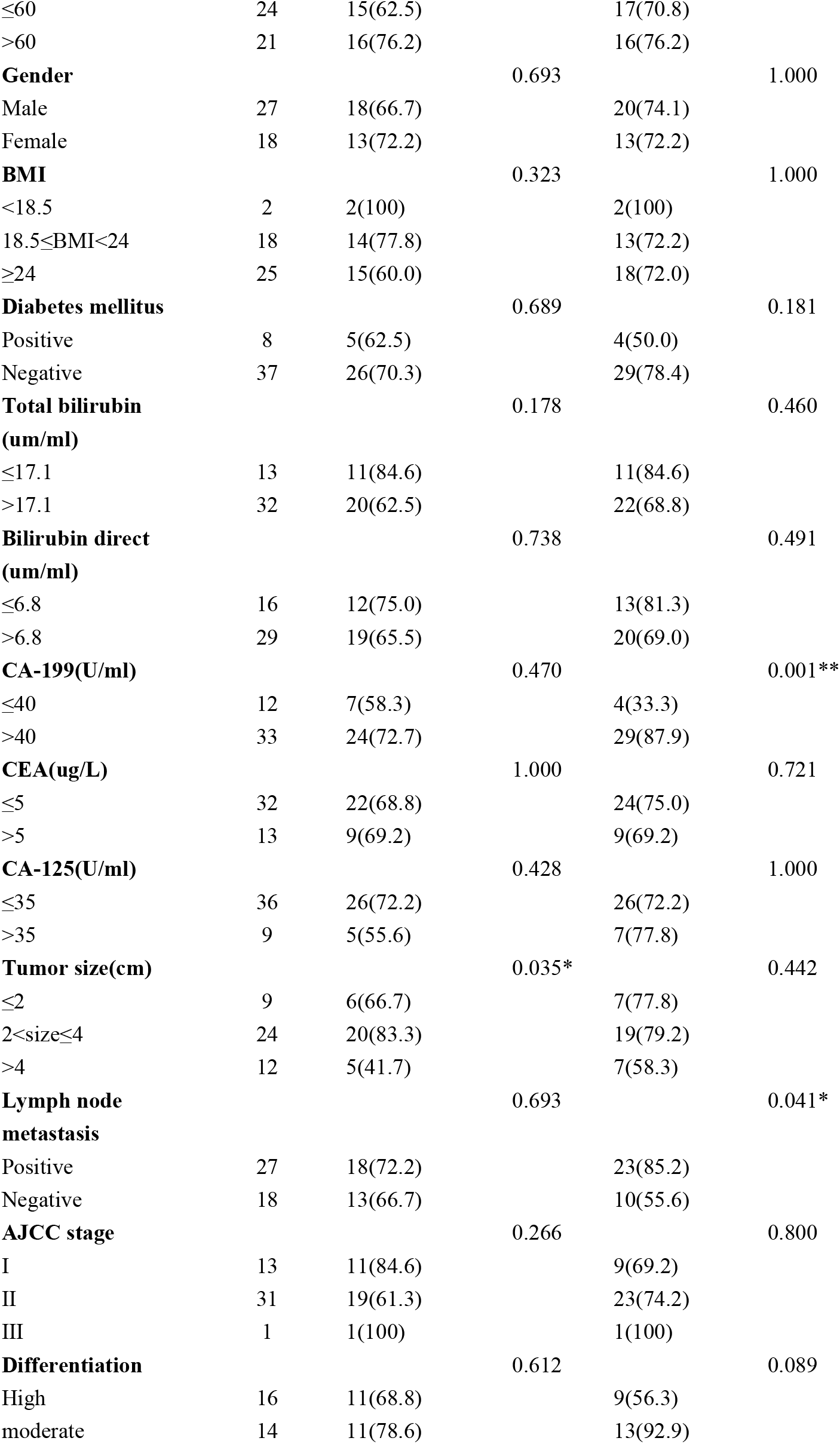
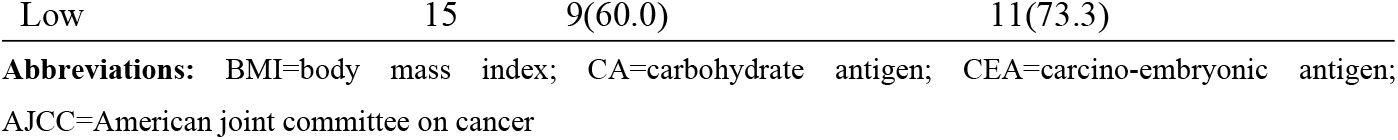
Relationship between the expression of β-catenin/LEF-1 and the clinical characteristics in PDAC patients.

**Table 4.**
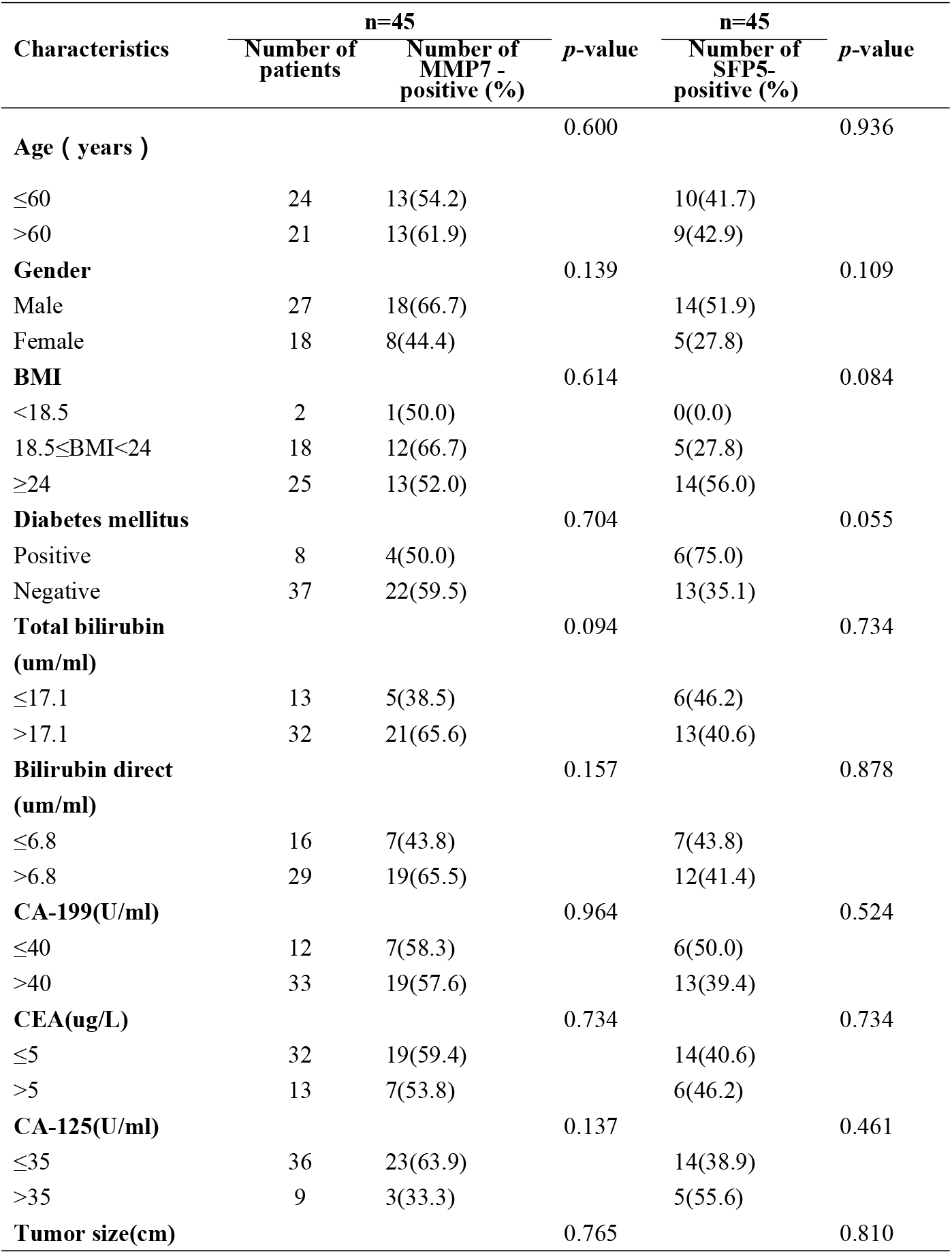
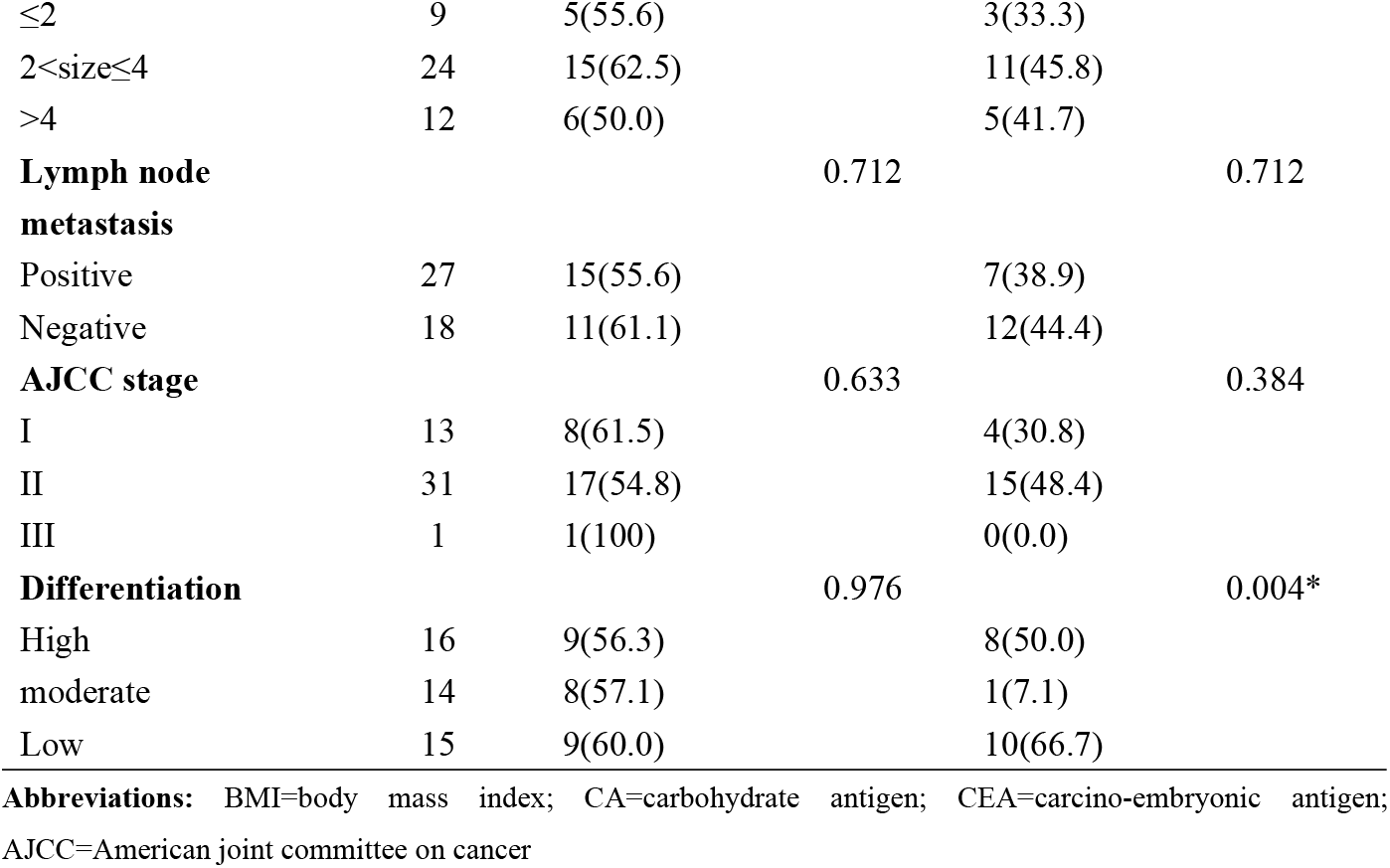
Relationship between the expression of MMP7/SFP5 and the clinical characteristics in PDAC patients.

### The long-term survival outcomes and influence factors of PDAC patients after radical resection

In this study cohort, the median survival times of all patients were 22 months (range:1 to 82months), The cumulative three-year and five-year OS rates were 7% and 2%, respectively (Figure 3A).

**Figure 3.**
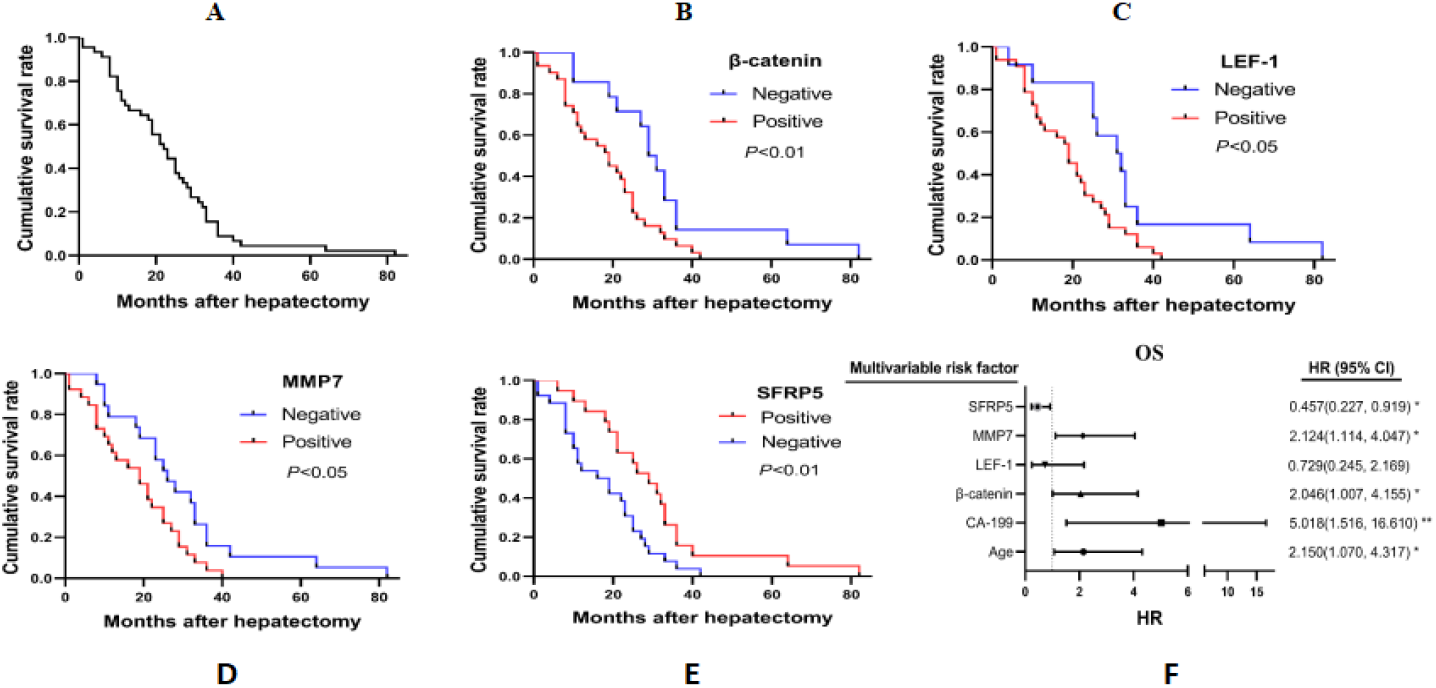
Cumulative survival curves and multivariate Cox regression analysis. **(A)** Cumulative survival curves in all patients. **(B)** Cumulative survival curves of patients in β-catenin positive and negative expression group. (C) Cumulative survival curves of patients in β-catenin positive and negative expression group. **(D)** Cumulative survival curves of patients in MMP7 positive and negative expression group. (E) Cumulative survival curves of patients in SFRP5 positive and negative expression group. (F) Forest plots of the results of multivariate Cox analysis for OS.

### β-catenin, LEF-1, MMP7 and SFRP5 positive expression were associated with prognosis in PDAC patients

The median survival times were 19.0 months, 19.0 months, 19.0 months and 29.0 months for the β-catenin, LEF-1, MMP7 and SFRP5 positive groups, while 29 months, 31 months, 26months and 17.5 months for the negative groups, respectively (*p*<0.05). This indicates that β-catenin, LEF-1 and MMP7 positive expression correlates with poor prognosis in PDAC patients, and SFRP5 positive expression correlates with longer survival times (Figure 3 B,C,D,E). To assess the factors related to survival in PDAC patients, univariate and multivariate analyses using the Cox proportional hazard regression model were performed (Table 4). As a result of multivariate analysis adjusting for the significant variables identified in the univariate analysis, age (*P*<0.05), CA-199 (*P*<0.01), β-catenin (*P*<0.05) and MMP7 (*P*<0.05) were identified as the independent risk factors for prognosis, while SFRP5 (*P*<0.05) was the independent protection factor (Figure 3F, Table 5).

**Table 5.** Univariate and multivariate Cox analysis for overall survival of PADC patients.

| Variables | OS |  |
| --- | --- | --- |
|  | Univariate | Multivariate |
|  | HR (95%CI) | HR (95%CI) |
| Age (≤60 years vs >60years) | 1.962(1.043, 3.692) * | 2.150(1.070, 4.317) * |
| Gender (male vs female) | 1.585(0.849, 2.958) |  |
| BMI (<18.5 vs 18.5≤BMI<24 vs ≥24) | 0.841(0.487, 1.453) |  |
| Diabetes mellitus (+ vs -) | 0.503(0.222, 1.137) |  |
| Total bilirubin (≤17.1um/ml vs >17.1 um/ml) | 0.580(0.295, 1.140) |  |
| Bilirubin direct (≤6.8um/ml vs >6.8 um/ml) | 0.615(0.321, 1.176) |  |
| CA-199(≤40 U/ml vs >40 U/ml) | 2.667(1.258, 5.656) * | 5.018(1.516, 16.610) ** |
| CEA (≤5 ug/L vs >5 ug/L) | 1.087(0.561, 2.104) |  |
| CA-125(≤35U/ml vs >35 U/ml) | 0.778(0.370, 1.637) |  |
| Tumor size (≤2cm vs 2<size≤4 vs >4cm) | 1.279(0.868, 1.883) |  |
| Lymph node metastasis (+ vs -) | 1.150(0.625, 2.114) |  |
| AJCC stage (I vs II vs III) | 1.108(0.576, 2.131) |  |
| Differentiation (High vs moderate vs Low) | 1.124(0.793, 1.593) |  |
| β-catenin (+ vs -) | 2.327(1.179, 4.593) * | 2.046(1.007, 4.155) * |
| LEF-1(+ vs -) | 2.253(1.097, 4.630) * | 0.729(0.245, 2.169) |
| MMP7(+ vs -) | 2.057(1.090, 3.881) * | 2.124(1.114, 4.047) * |
| SFRP5(+ vs -) | 0.465(0.249, 0.870) * | 0.457(0.227, 0.919) * |

\* $P<0.05$ , \*\* $P<0.01$ , \*\*\* $P<0.001$ .

### The establishment and evaluation of the line graph model

R software was used to establish a rosette model to predict the prognosis of pancreatic cancer according to the results of multi-factor analysis. According to the column chart, age ≤60 is 0 points;Age >60, the score is 57.5; CA199≤40 was 0, CA199>40 is 100 points; β-catenin negative score was 0, β-catenin positive score was 50; MMP7 negative score was 0, positive score was 55; SFRP5 positive score was 0, negative score was 57.5. The higher the total score of the column chart, the lower the corresponding OS in 3 years and 5 years (Figure. 4). The differentiation test results indicate that the c-index of the line graph model is 0.742, indicating a high accuracy of the model. By drawing the calibration diagram of predicted value and actual value, consistency test was carried out. The 3-year and 5-year OS predicted by the rosette model had a good correlation with the actual 3-year and 5-year OS (Figure. 5).

**Figure 4.**
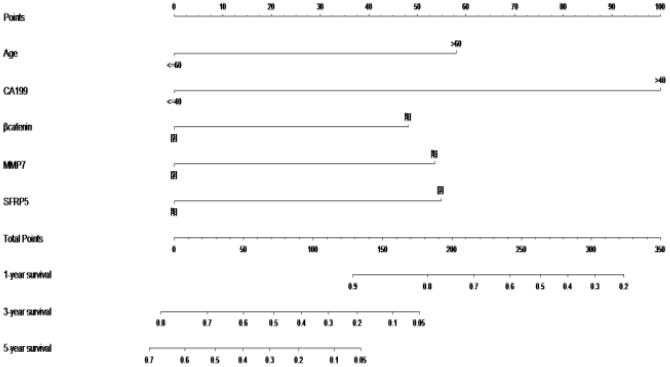
In the prediction model of the prognosis of pancreatic cancer, the score of each factor was obtained according to the upper scale, and the total score was obtained by adding the scores of each factor. From the total

**Figure 5.**
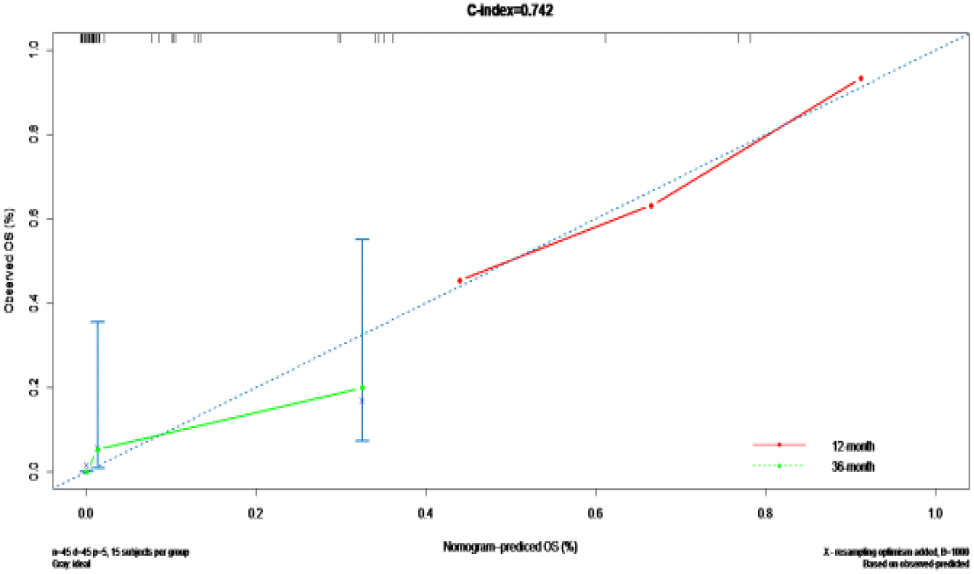
Compare the calibrated plots of 3- and 5-year overall survival predicted by the rotigrams with the observed 3- and 5-year overall survival.

## Discussions

Wnt signalling continues to play indispensable roles in tissue homeostasis, cell renewal, and regeneration[13]. It has been wildly reported in pancreatic cancer. SMARCAD1 activates Wnt/β-catenin signaling pathway by altering the expression of β-catenin to induces EMT and promotes pancreatic cancer metastasis[14]. USP21 overexpression and its nuclear localization correlate positively with PDAC, USP21 interacts with, deubiquitinates and stabilizes the Wnt pathway transcription factor TCF7 to activate gene expression in the Wnt network^[^15]. Following Wnt activation, β-catenin can translocate from cytokines to the nucleus, where β-catenin binds to the T-cell factor/lymphoid enhancer-binding factor (TCF/LEF) to regulate the expression of Wnt target genes, thus playing an important role[16, 17]. LEF-1 is one member of the TCF-LEF subfamily and belong to a family of high-mobility-group proteins that work as a downstream transcription factor of β-catenin signaling. Moreover, one of the unique features of LEF-1 is to regulate target gene expression by DNA bending and helical phasing of transcription-binding sites, acting as an “architectural” transcription factor[18]. Evidence has shown that expression of LEF-1 was increased in multiple malignant diseases including hepatocellular carcinoma, acute myeloid leukaemia, malignant melanoma, pancreatic carcinoma[19-22]. Matrix metalloproteinase-7 (MMP-7), a smallest secreted proteolytic enzyme with broad substrate specificity against ECM components, is one of the important downstream target genes of β-catenin/TCF-4[23, 24]. Secreted frizzled-related protein (SFRP), an evolutionary conserved family of secreted proteins, may block Wnt signaling either by binding to Fz proteins or by forming nonfunctional complexes with Fz[25]. SFRPs represent 5 types of proteins: SFRP1–SFRP5[26]. there are many regulatory factors, so Wnt signaling is a vital molecular pathway in pancreatic cancer and may be amenable to targeted drug therapy [27].

In our study, the median survival times of 45 PADC patients after radical pancreatoduodenectomy were 22 months, the cumulative three-year and five-year OS rates were 7% and 2% respectively, and were significantly lower (35.6% and 31.7%) compared with a study by Maeda S et al. (including 45 patients who had total pancreatectomy from 2007 to 2016)[28], this may be related to difference in time, region, equipment, or technology. It can be sure that the prognosis of PDAC remain dismal, despite underwent radical pancreatoduodenectomy. It necessitates the urgent hunt for novel more effective therapeutic strategies.

Our study also found that in pancreatic cancer, β-catenin, LEF-1 and MMP7 genes showed not only high mRNA expression, protein but also showed higher positive rates which were associated with adverse pathological characteristics. Patients in the β-catenin positive expression group exhibited a bigger tumor size than those in the negative expression group. Patients with positive LEF-1 showed a higher serum CA19-9 level and positive lymph node metastasis. Previous studies found that LEF-1 is essential to T cell growth and development, the high expression of LEF-1 is closely associated with several types of malignancies, including leukemia, lymphoma[29, 30], it may promote lymph node metastasis. SFRP5 expression was related to tumor differentiation. Our study suggests that Wnt signaling pathways and its four key regulators may be involved in the occurrence and development of PDAC.

The patients with positive expression of β-catenin, LEF-1 and MMP7 had a shorter survival time than the patients with negative expression in PDAC. while SFRP5 was contrary to them and negatively correlated with LEF-1 protein expression. In addition, Age, CA-199, β-catenin, MMP7 and SFRP5 could serve as promising biomarkers for predicting prognosis in PDAC patients, however, LEF-1 expression does not seem to be an independent prognostic factor in PDAC. CA-199, the most extensively used biomarker in pancreatic cancer, can be used to evaluate staging, prognosis, tumor resectability, recurrence and treatment effect, but it may be falsely positive in cases of biliary infection, inflammation, or obstruction (regardless of aetiology) and has certain limitations[31, 32]. The efficacy of multi-indicator joint assessment needs to be further validated in clinical practice. In this study, we established a rograph prediction model for pancreatic cancer based on 5 independent prognostic factors. The results suggested that 0 points were scored when age ≤60, and 57.5 points were scored when age >60. CA199≤40 was 0 points, CA199>40 was 100 points; β-catenin negative score was 0, β-catenin positive score was 50; MMP7 negative score was 0, positive score was 55; SFRP5 positive score was 0, negative score was 57.5. By calculating the scores of age, CA199, β-catenin, MMP7 and SFRP5 by the rograph, we can intuitively assess the prognosis of patients.

Taken together, Wnt signaling pathways can be a potential therapeutic target , of which β-catenin , MMP7, SFRP5, Age and CA199 could serve as a promising biomarker for predicting prognosis in PDAC patients. Furthermore, the lipophrograph prediction model based on these factors is established. The rosette prediction model helps to obtain prognostic estimates at the individual level. At the same time, the unique visualization effect of the line map helps surgeons intuitively understand the contribution weight of each level of each prognostic factor to the long-term survival of patients. Based on this model, it is helpful for clinicians to screen patients with poor prognosis for closer follow-up or adjuvant treatment.

## Human subjects research

The study was approved by the Independent Ethics Committee of Tumor Hospital Affiliated to Xinjiang Medical University in 2020(G-2020020). All patients signed the informed consent form and agreed to participate in the study.

## Consent for publication

All the listed authors carefully examined and approved this manuscript. We all agree to submit it to [PLOS One] for publication and understand and accept the publication policy of this journal.

## Availability of data and material

Informed consent was obtained from all individual participants who were included in the study. The data supporting the conclusions of this article are included within the article. For the present study, the raw datasets used and analysis are available from the corresponding author upon reasonable request.

## Authors’ contributions

Panpan Kong was responsible for experimental design and article writing. Feng He was responsible for the analysis and processing of experimental data. Huifang Liu was responsible for collecting clinical data and conducting bioinformatics analysis. Dong Yan was responsible for revising, polishing, and providing financial support for the article.

## Acknowledgements

The author would like to express gratitude to all those who have contributed to the research and writing of this paper. Special thanks are extended to Professor Dong Yan, the corresponding author, for providing financial support and assistance in designing the article.

